# A collection of split-Gal4 drivers targeting conserved signaling ligands in *Drosophila*

**DOI:** 10.1101/2024.10.10.617664

**Authors:** Ben Ewen-Campen, Neha Joshi, Ashley Suraj Hermon, Tanuj Thakkar, Jonathan Zirin, Norbert Perrimon

**Affiliations:** Department of Genetics, Blavatnik Institute, Harvard Medical School, Boston, MA 02115, USA; Howard Hughes Medical Institute, Boston, MA 02115, USA

## Abstract

Communication between cells in metazoan organisms is mediated by a remarkably small number of highly conserved signaling pathways. Given the relatively small number of signaling pathways, the existence of multiple related ligands for many of these pathways is thought to represent a key evolutionary innovation for encoding complexity into cell-cell signaling. Relatedly, crosstalk and other interactions between pathways is another critical feature which allows a modest number pathways to ultimately generate an enormously diverse range of outcomes. It would thus be useful to have genetic tools to identify and manipulate not only those cells which express a given signaling ligand, but also those cells that specifically co-express pairs of signaling ligands. Here, we present a collection of split-Gal4 knock-in lines targeting many of the ligands for highly conserved signaling pathways in *Drosophila* (Notch, Hedgehog, FGF, EGF, TGF*β*, JAK/STAT, JNK, and PVR). We demonstrate that these lines faithfully recapitulate the endogenous expression pattern of their targets, and that they can be used to specifically identify the cells and tissues that co-express pairs of signaling ligands. As a proof of principle, we demonstrate that the 4th chromosome TGF*β* ligands *myoglianin* and *maverick* are broadly co-expressed in muscles and other tissues of both larva and adults, and that the JAK/STAT ligands *upd2* and *upd3* are partially co-expressed from cells of the midgut following gut damage. Together with our previously collection of split-Gal4 lines targeting the seven Wnt ligands, this resource allows *Drosophila* researchers to identify and genetically manipulate cells that specifically express pairs of conserved ligands from nearly all the major intercellular signaling pathways.

## Introduction

Coordinated activity between cells in multicellular organisms requires the activity of intercellular signaling pathways, where signaling cells generate ligands that are received and interpreted by cells which express relevant receptors. A relatively small number of intercellular signaling pathways are extraordinarily highly conserved across animal evolution; these pathways include Notch, Hedgehog, fibroblast growth factor (FGF), epidermal growth factor (EGF), Wnt/Wingless, transforming growth factor *β* (TGF*β*), Janus kinase/signal transducer and activator of transcription (JAK/STAT), Jun kinase (JNK), platelet-derived growth factor/vascular endothelial growth factor-related receptor (PVR), and others (Perrimon et al. 2012; Housden and Perrimon 2014).

Despite the relatively small number of highly conserved signaling pathways, an enormous complexity of outcomes is possible. Among the many features that enable complexity in metazoan signaling is the fact that there are often multiple closely related ligands capable of activating a given pathway (Perrimon et al. 2012; Housden and Perrimon 2014). In the *Drosophila* genome, for example, while there is only a single characterized ligand for the Hedgehog pathway (*hedgehog [hh]*) and for the JNK pathway (*eiger [egr]*), there are two paralogous ligands for the Notch receptor (*Delta* and *Serrate*), three FGF ligands (*bnl, ths, pyr*), three JAK/STAT ligands (*upd1*-3), three PVR ligands (*Pvf1-3*), four EGF agonistic ligands (*spitz, gurken, Keren*, and *vein*), and seven ligands each for Wnt and TGF*β* signaling (Nusse 2001; Upadhyay et al. 2017). The fact that, in many cases, these ligand families expanded and diversified early in animal evolution and have been very highly conserved ever since suggests that ligand diversity is a critical feature which helps to allow this modest number of signaling pathways to exhibit the enormous range of outputs that is observed *in vivo*.

Paralogous ligands can have largely overlapping functions, wholly independent functions, or a complex combination of the two (Ewen-Campen et al. 2017). For example, in the *Drosophila* midgut, the JAK/STAT ligand *upd1* plays a critical function in intestinal stem cells (ISCs), where it promotes proliferation under basal conditions; neither *upd2* nor *upd3* is required for this function (Osman et al. 2012). However, when the midgut epithelium is damaged, *upd2* and *upd3* are upregulated in enterocytes (ECs), and function in an additive manner to promote damage-responsive ISC proliferation (Osman et al. 2012). Importantly, while *upd2* and *upd3* function together in the context of midgut regeneration, in other contexts the three *upd* ligands have wholly separate functions, such as the role that *upd2* plays in the fat body to regulate systemic metabolism (Rajan and Perrimon 2012).

In order to tease apart the function of a pair of related genes, it is often necessary to understand their expression and co-expression patterns in space and time. The split-Gal4 system is particularly well suited to identify those cells in which two genes are co-expressed (Luan et al. 2006; Luan et al. 2020). In this system, a transcriptional activator domain (either VP16 or p65) and a DNA-binding domain (from Gal4), each separately fused to a leucine zipper, are separately expressed under the control of two different regulatory sequences. In those cells that express both components, and only in those cells, the leucine zippers bind to another to constitute a functional transcription factor that activates UAS-based transgenes (Luan et al. 2006; Luan et al. 2020). Thus, the split-Gal4 system can be used to identify *in vivo* those cells that co-express two genes of interest.

The split-Gal4 system has predominantly been used to generate lines driven by minimal enhancer fragments, in order to specifically label very small numbers of neurons in the brain (Tirian and Dickson 2017). However, recent studies have shown that the split-Gal4 components can also be inserted directly into a coding sequence of a gene of interest, either downstream of a T2A sequence and/or encoded in a “trojan exon”, which allows for split-Gal4 lines that recapitulate the entire expression pattern of a gene-of-interest (Ewen-Campen et al. 2020; Chen et al. 2023; Ewen-Campen et al. 2023; Diao et al. 2024).

Previously, we generated a collection of knock-in split-Gal4 lines targeting the seven Wnt ligands, and demonstrated how these lines can be used to identify cells and tissues which co-express certain pairs of Wnt agonists *in vivo* (Ewen-Campen et al. 2020). Here, we present an expanded collection of knock-in split-Gal4 lines targeting the ligands from additional highly conserved signaling pathways in *Drosophila*: Notch, Hedgehog, FGF, EGF, TGF*β*, JAK/STAT, JNK, and PVR. These lines can be used to identify tissues that co-express pairs of ligands-of-interest, and more generally may be useful to researchers wishing to genetically manipulate specific cell populations that can be intersectionality-labeled using these reagents. Furthermore, these reagents can be used to identify crosstalk between different pathways by identifying cells that co-express ligands for more than one pathway.

## Results and Discussion

### A resource of split-Gal4 knock-in lines targeting genes encoding conserved signaling ligands

We created CRISPR knock-in donor plasmids to insert either T2A-p65 or T2A-Gal4DBD into each of the ligands shown in Table 1. We designed these donor constructs to be inserted in a coding exon (Figure 1A), targeting shared exons for any target gene with multiple isoforms. These inserts target coding exons, and therefore are presumed to produce loss-of-function alleles of the target gene. These knock-in constructs also contain a 3xP3-RFP fluorescent marker used to recover transformants, and this marker is flanked by loxP sites so that it can be removed in the future by crossing to a line containing a source of *Cre*. When a p65 knock-in line is crossed to a Gal4DBD knock-in line, those cells that co-express both ligands can be visualized using a UAS-based reporter and this population of cells can be genetically manipulated (Figure 1B,C).

**Table 1.**
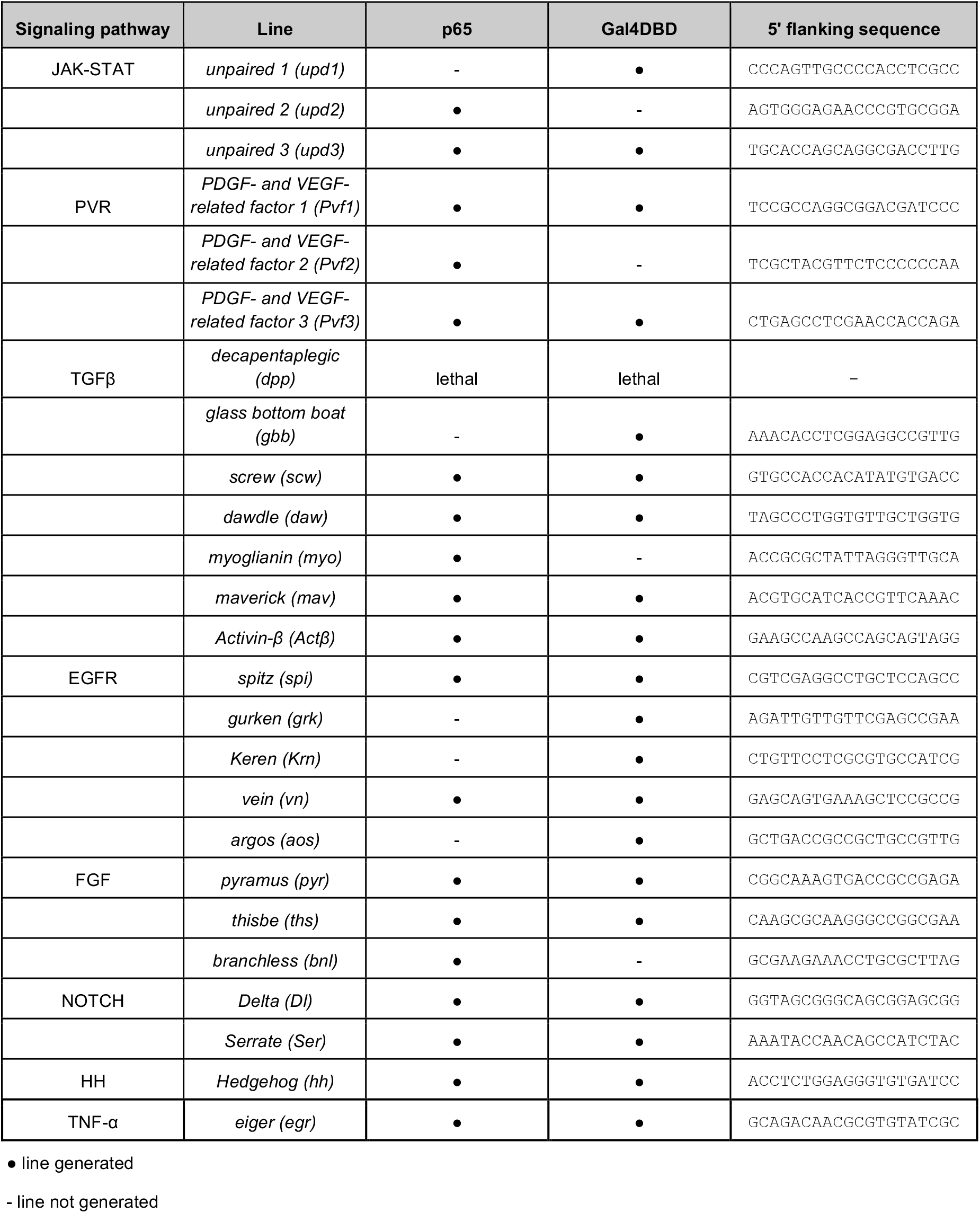
Lines generated in this study.

**Figure 1.**
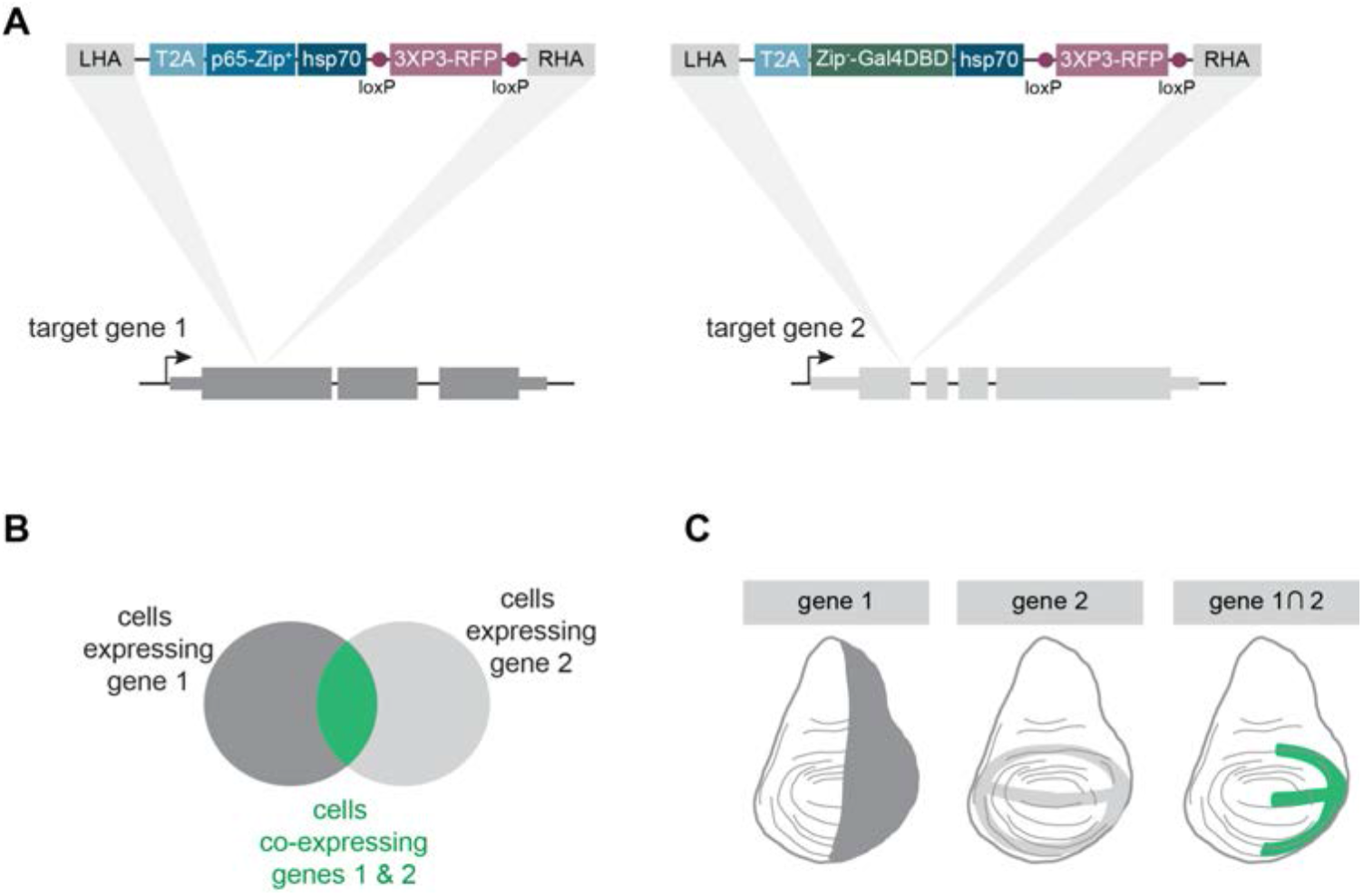
Strategy for creating split-Gal4 knock-in lines for conserved signaling ligands. (A) Knock-in cassettes containing T2A-p65 or T2A-Gal4DBD were inserted using CRISPR-Cas homology-directed repair into a coding exon of each signaling ligand, so that the expression domain of each split-Gal4 line captures the native expression pattern of the target gene. LHA = left homology arm. RHA = right homology arm. 3xP3-RFP = fluorescent marker for transgenesis. (B) Schematic of the split-Gal4 system. Only in those cells which co-express both target genes are the two split-Gal4 components reconstituted via leucine-zipper binding into a functional transcription factor. (C) Cartoon example of two hypothetical split-Gal4 lines in a developing wing disc, demonstrating how the expression of a UAS-based transgene is limited to only those cells that co-express both gene 1 and gene 2. Dorsal is up, anterior is left.

For each target gene, we first confirmed that the insertion segregates on the expected chromosome, and then validated that the insertion was correct using PCR-based genotyping and sequencing, recapitulation of a known *UAS:GFP* expression pattern, or both.

We successfully recovered 39 transgenic lines: both p65 and Gal4DBD for 15 genes (totaling 30 lines), and either p65 or Gal4DBD knock-ins for 9 additional genes (Table 1). We were unable to recover any knock-ins for *dpp*, despite multiple attempts at the extreme 3’ end of the coding sequence, which is consistent with the fact that this gene is a dominant lethal (Wharton et al. 1993)(Table 1). Altogether, this collection should allow for analysis of nearly all pairwise combinations of ligands within a signaling family, and many combinations across pathways.

To determine whether our knock-in lines successfully recapitulate the expression pattern of the genes into which they are inserted, and whether the T2A-p65 and T2A-Gal4DBD lines mirror one another for a given target, we examined their expression patterns in L3 larval wing discs. We crossed each of our knock-in lines to a reciprocal “tester” line, containing either *tub:VP16-AD* or *tub:Gal4DBD* and a *UAS:2xEGFP* reporter. We focused on those genes for which we obtained both p65 and Gal4DBD lines, and those genes predicted to be expressed in a specific pattern in wing discs.

The split-Gal4 lines generated in this study successfully capture the endogenous expression pattern of their target genes, and we generally observed that p65 lines and Gal4DBD for a given target gene were essentially congruent (Figure 2). We found that both p65 and Gal4DBD lines targeting *hh* drive expression in the posterior compartment of the wing disc (Lee et al. 1992; Tabata et al. 1992)(Figure 2), and that lines targeting the Notch ligands *Delta* and *Ser* recapitulate well-characterized expression patterns (Doherty et al. 1996; Yan et al. 2003). Lines targeting the FGF ligands *pyr* and *ths* are both expressed in the epithelial cells of the notum, overlaying the adult muscle precursors (AMPs), as previously reported (Everetts et al. 2021; Patel et al. 2022) (Figure 2). *vn*-*p65/Gal4DBD* lines mirror endogenous patterns of *vn* expression (Simcox et al. 1996).

**Figure 2.**
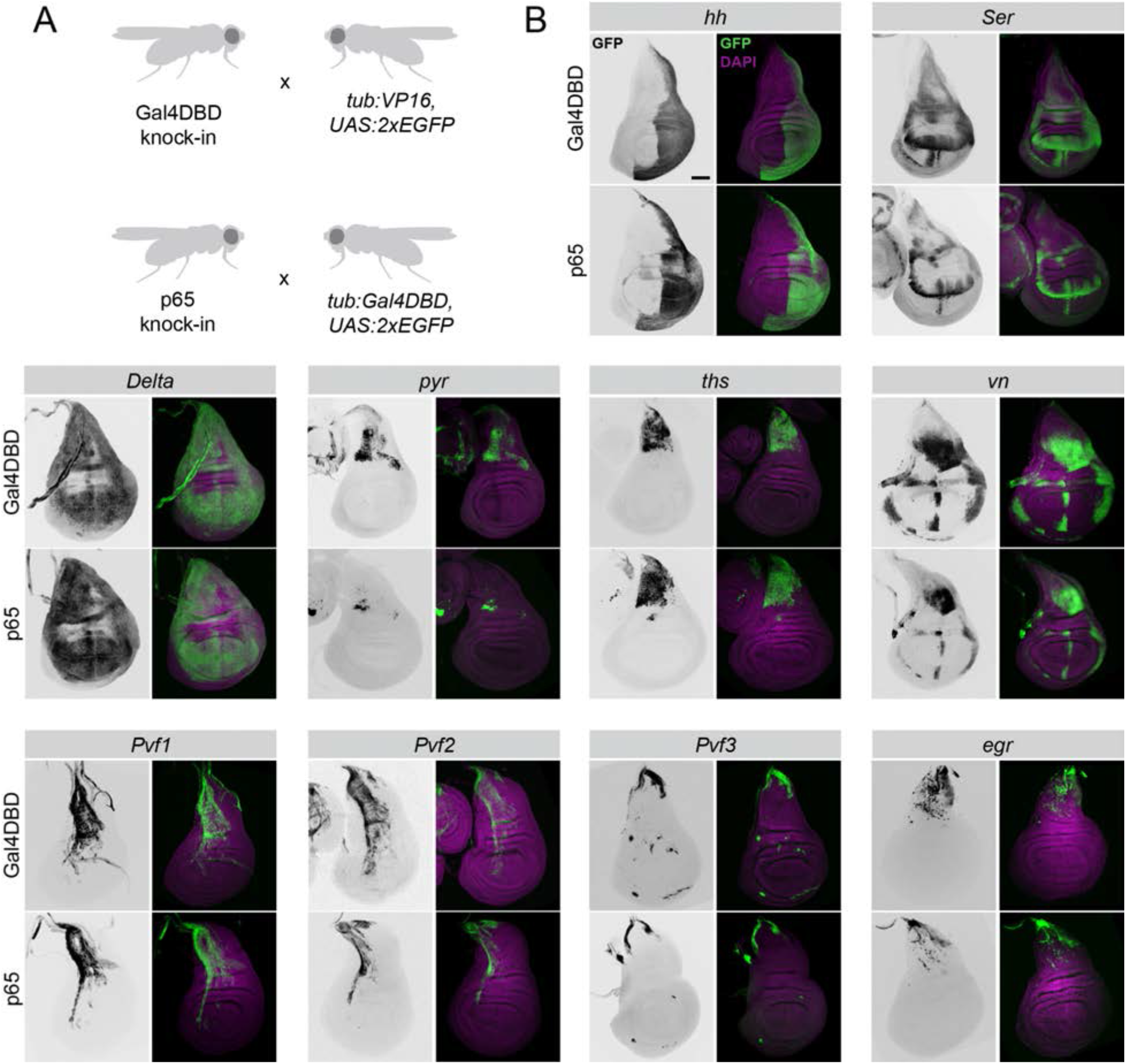
Expression patterns for a selection of split-Gal4 lines in the L3 wing disc. (A) The full expression pattern of each knock-in line was determined by crossing to a ubiquitously-expressed split-Gal4 “tester line”: *tub:VP16, UAS:2xEGFP* for Gal4DBD knock-ins and *tub:Gal4DBD, UAS:2xEGFP* for p65 knock-ins. (B) For each target gene, transgenic lines expressing Gal4DBD (top row) or p65 (bottom row) are shown. Dorsal is up, anterior is left.

The expression patterns of other ligands are not well characterized in the wing disc, but several lines of evidence suggest that the expression patterns we observe mirror endogenous gene expression patterns. For example, to our knowledge the expression patterns of *Pvf1, Pvf2*, and *Pvf3* have not been characterized in the wing disc, but we observed that *Pvf1* and *Pvf2* expression were restricted to the apical structures of the disc, as described for *PVF1* protein (Rosin et al. 2004), and enriched primarily in the notum, and that *Pvf3* expression was primarily found at the dorsal edge of the notum and the stalk, in a location where the *PVR* receptor is known to function in dorsal closure (Ishimaru et al. 2004). These expression patterns were also consistent with *in silico* predictions from single cell RNAseq (scRNAseq) studies of the wing disc (Deng et al. 2019; Worley et al. 2022). Relatedly, the JNK ligand *egr* is also enriched near the stalk and notum, where the JNK target gene *puc* is endogenously expressed (Agnes et al. 1999), and where *in silico* predictions indicate *egr* is enriched (Deng et al. 2019).

While the expression patterns between p65 and Gal4DBD lines were largely similar, we observed that in some cases, the expression of our Gal4DBD lines were notably stronger than the p65 lines targeting the same ligands (Figure 2, see e.g. *Ser, pyr, vn*). We noted that these differences could be caused by differential strengths of the two “tester” lines we used; specifically, the *tub:Gal4DBD, UAS:2xEGFP* tester line may drive weaker expression than the *tub:VP16-AD, UAS:2xEGFP* line.

To test whether this was the case, we crossed a variety of our knock-in split-Gal4 lines to one another, in both directions. We examined expression of *hh-p65 ∩ Ser-Gal4DBD*, and *hh-Gal4DBD ∩ Ser-p65*, and performed similar bidirectional crosses between *vn ∩ Ser*. In each case, we observed essentially indistinguishable expression patterns regardless of which gene was driving p65 versus Gal4DBD, indicating that these lines drive very similar expression levels to one another (Figure 3), and indicating that the *tub:VP16-AD, 2xEGFP* tester line likely drives lower expression than the *tub:Gal4DBD, UAS:2xEGFP* line. Altogether, these results demonstrate that these knock-in split-Gal4 lines successfully recapitulate the expression patterns of their target genes, and that the expression levels of p65 and Gal4DBD knock-in lines are very similar to one another.

**Figure 3.**
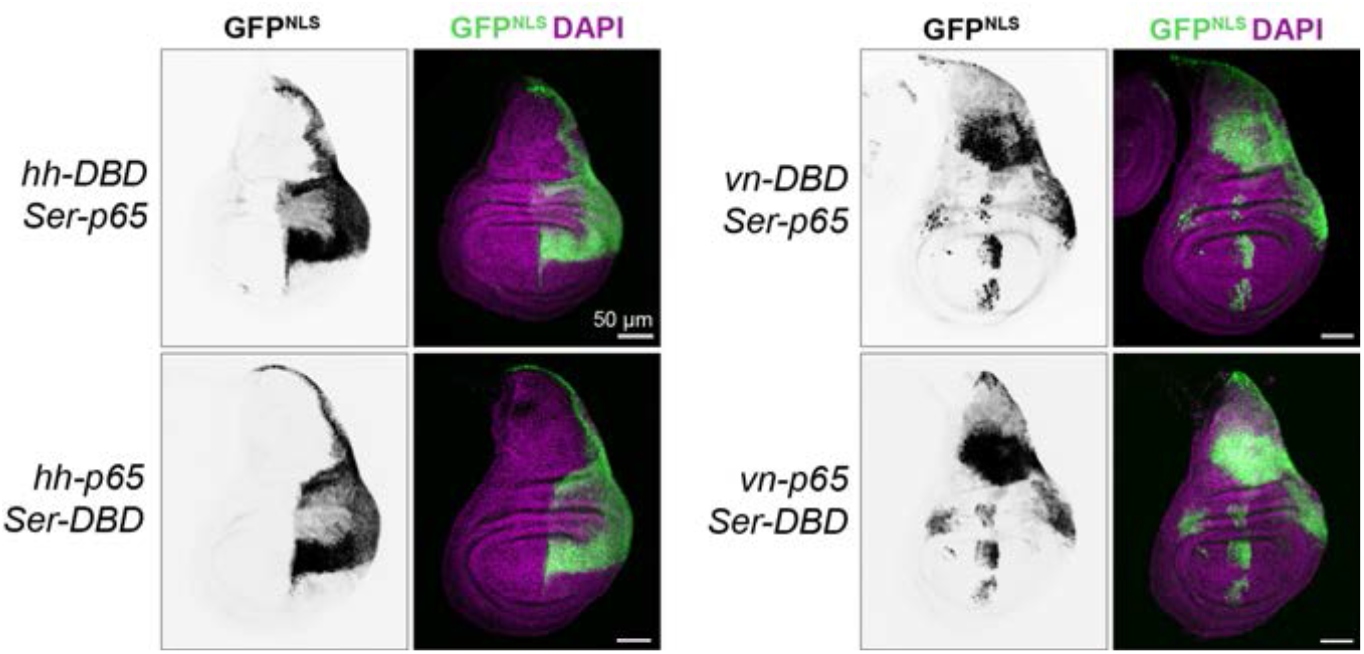
Intersectional labeling by split-Gal4 lines. Expression patterns in the L3 wing disc of *hh ∩ Ser* and *vn ∩ Ser*, in each possible p65/Gal4DBD configuration. Nuclear-localized UAS:GFP^Stinger^ was used in these experiments. Dorsal is up, anterior is left.

### TGF*β* ligands *maverick* and *myoglianin* are co-expressed broadly in larval and adult muscles

To demonstrate possible applications for this collection of split-Gal4 lines, we first turned to the seven TGF*β* family ligands. Using the bioinformatic tool Paralog Explorer (https://www.flyrnai.org/tools/paralogs/web/)(Hu et al. 2022), we queried the co-expression correlation of each of the seven TGF*β* ligands against the others across four high throughput RNAseq datasets from the modENCODE project: across tissues, across cell lines, across developmental datasets, and across experimental treatments. This analysis suggested that *maverick (mav)* and *myoglianin (myo)* are consistently among the TGF*β* ligands with the most highly correlated co-expression across all four categories of comparison.

*myo* and *mav* are both located on the 4th chromosome. Along with *Actβ* and *daw, myo* is a member of the Activin family of ligands (Lo and Frasch 1999; Upadhyay et al. 2017), and is orthologous to mammalian GDF11 and Myostatin (Demontis et al. 2014). In contrast, *mav* does not fall neatly into either the Activin or BMP families (Nguyen et al. 2000; Upadhyay et al. 2017), and phylogenetic analysis of TGF*β* ligands has not definitively identified which homolog is most closely related to *mav* (Zee et al. 2008). Loss-of-function *myo* mutants are pupal lethal, and this ligand has been shown to be expressed in both glia and in muscles where it plays a number of critical signaling roles (Awasaki et al. 2011; Demontis et al. 2014; Augustin et al. 2017; Upadhyay et al. 2020). *mav* loss-of-function mutants are viable, but display neurological defects (Myers et al. 2017; Hoyer et al. 2018).

Given the high degree of *myo* and *mav* co-expression predicted *in silico*, we examined their co-expression *in vivo*. In L3 larva, we found that *myo-p65* was expressed broadly in all muscles of the body wall, as well as detectable in oenocytes and trachea (Figure 4A). *mav-Gal4DBD* was detected in larval muscles as well as trachea (Figure 4A), and intersection labeling of *myo ∩ mav* was detected in all larval muscles as well as in trachea (Figure 4A). In adults, *myo ∩ mav* co-expression was again detected in all muscles of the body wall and the legs, but largely excluded from the flight muscles of the thorax (Figure 4B). As in larva, *myo-p65* expression was broader than *mav-Gal4DBD*, and could be detected in fat body, salivary gland and other tissues, whereas *mav-Gal4DBD* appeared largely restricted to the muscles and trachea (Figure 4B.) Taken together, these results suggest that *myo* and *mav* are broadly co-expressed in both muscles and trachea throughout larval and adult life, and that future studies may benefit from double knock-out experiments to examine the possibility of overlapping function.

**Figure 4.**
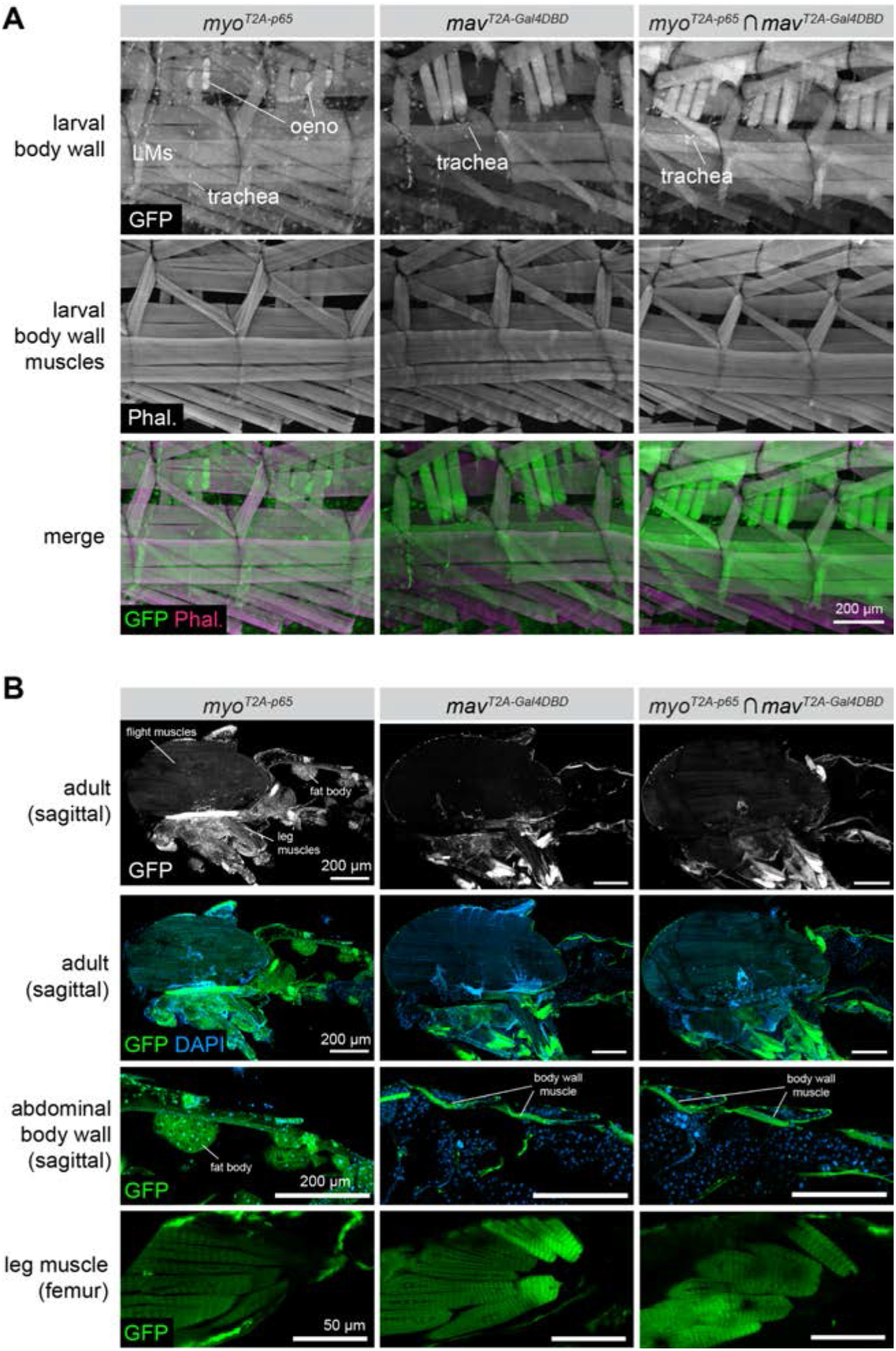
The TGF*β* ligands *maverick* and *myoglianin* are co-expressed in muscles of the larva and adult. **(A)** Expression of *myo-p65, mav-Gal4DBD*, and co-expression of *myo-p65 ∩ mav-Gal4DBD* in L3 larval body wall tissues, including muscles, trachea, and oenocytes. Both ligands are expressed broadly in nearly all larval body-wall muscles. LM = larval muscles; Phal = phalloidin; oeno = oenocytes. **(B)** Expression of *myo-p65, mav-Gal4DBD*, and *myo-p65 ∩ mav-Gal4DBD* in sagitally sectioned adult carcass (top three rows) and in adult leg muscles (bottom row).

As a second example of how this collection of split-Gal4 lines can be used, we examined the co-expression of JAK/STAT ligands in the adult gut, where all three *upd* ligands play important functions (Jiang et al. 2009; Osman et al. 2012). All three ligands are upregulated in the midgut in response to tissue damage, with *upd3* showing the highest levels of damage-responsive upregulation (Jiang et al. 2009; Osman et al. 2012), and it has been shown previously that *upd2* and *upd3* have an additive effect on gut response to damage, with *upd2, upd3* double mutants showing stronger effects than either of the single mutants (Osman et al. 2012). However, it is known whether the same cells in the gut upregulate *upd2* and *upd3*, or whether they are upregulated in distinct gut cell populations.

We first examined the expression of *upd2-p65* and *upd3-Gal4DBD* in control midguts and in midguts of flies which had been damaged by bleomycin feeding. As expected, both *upd2* and *upd3* were strongly upregulated by bleomycin-induced damage (Figure 5), with *upd3* showing dramatically stronger upregulation (Figure 5). When we examined the intersectional staining of *upd2 ∩ upd3*, we observed a notable increase in the population of GFP+ cells in the midgut, indicating that many individual cells in the midgut simultaneously upregulate the expression of *upd2* and *upd3* in response to damage, whereas many other cells must upregulate *upd3* alone given the differential expression pattern of *upd3-Gal4DBD* versus *upd2 ∩ upd3*. Altogether, these experiments demonstrate that our collection of split-Gal4 lines faithfully recapitulates dynamic expression changes in response to tissue damage, and can additionally be used to identify those cells that specifically upregulate pairs of genes simultaneously.

**Figure 5.**
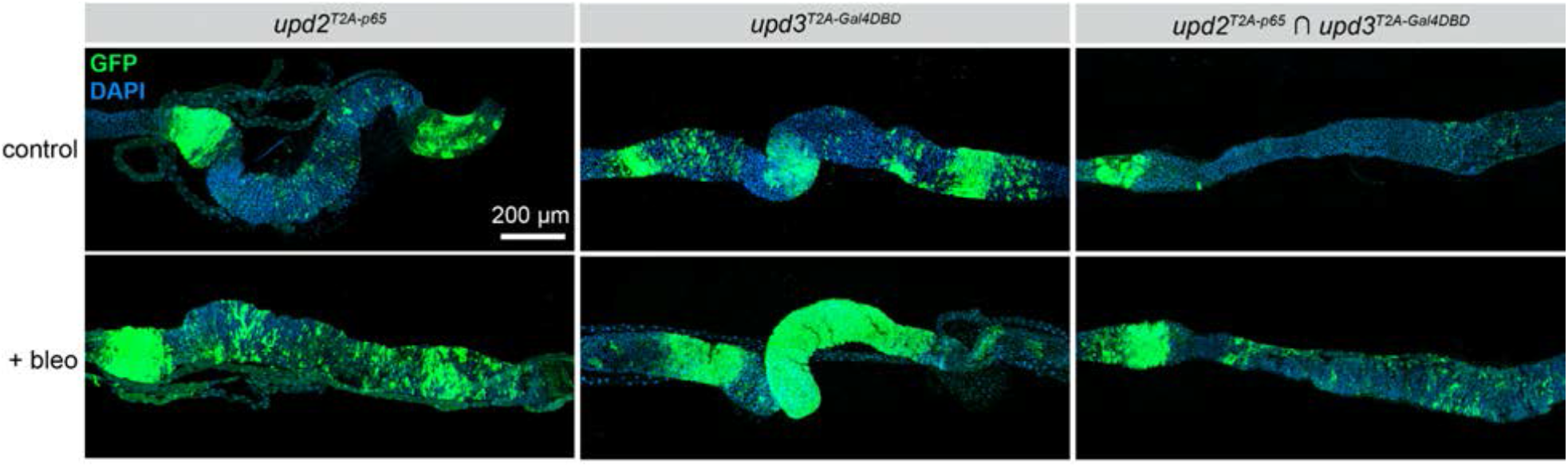
The JAK/STAT ligands *upd2* and *upd3* are co-expressed in cells of the posterior midgut in homeostasis and in response to cellular damage. Expression of *upd2-p65, upd3-Gal4DBD*, and co-expression of *upd2-p65 ∩ upd3-Gal4DBD* in the adult posterior midgut in control guts (top row) and guts damaged by bleomycin treatment (bottom row). Anterior is to the left.

Previously, identifying the cells or tissues that co-express a pair of genes-of-interest required such techniques as double antibody staining or double *in situ* hybridization. The creation of intersectional genetic labeling systems such as split-Gal4 system provided a genetic means to specifically label such cells, and has the additional benefit that these cells can be genetically manipulated in addition to simply being labeled. Here, we describe a collection of split-Gal4 lines targeting a collection of highly conserved signaling ligands, many of which belong to closely related paralogs. We demonstrate that these lines recapitulate the endogenous expression patterns, and that they can be used to identify cells or tissues where genes-of-interest are co-expressed.

The resource described here, together with our previous collection of knock-in split-Gal4 lines targeting the Wnt family of ligands (Ewen-Campen 2020), will allow detailed analysis of most major signaling pathway ligand expression and co-expression in *Drosophila*. We believe these reagents will be useful to others in the field, and have made them available at the Bloomington Stock Center.

We also note that these reagents may be useful for the annotation of scRNAseq datasets. As we and others have previously shown, split-Gal4 can be a powerful technique to map *in silico* cluster predictions to *in vivo* anatomy because most clusters cannot be uniquely identified using a single marker gene (Chen et al. 2023; Ewen-Campen et al. 2023). Given the widespread expression of many signaling ligands in multiple tissues throughout development and in the adult, the lines described here may prove useful tools to locate and study cell-types first identified via scRNAseq.

## Methods and Materials

### Experimental Animals

*Drosophila melanogaster* stocks were maintained and crossed on standard laboratory cornmeal food, with experiments performed at 25ºC. “Split-Gal4 tester lines” and UAS:2xEGFP lines were obtained from the Bloomington Drosophila Stock Center: *tub:Gal4DBD* ; *UAS:2xEGFP* (BL60298) and *tub:VP16[AD], UAS:2xEGFP* (BL60295), and UAS-2xEGFP, *w* ; *Sp / CyO* ; *UAS-2XEGFP* (BL60293), *w* ; *UAS-2XEGFP* ; *Dr / TM3,Sb* (BL60292); and *w* ; *UAS-Stinger* ; *MKRS/TM6b* (BL90920).

### Cloning of knock-in constructs

Coding sequences for p65-Zip^+^ and Zip^-^-Gal4DBD sequences were originally amplified from pBPp65ADZpUw (Addgene 26234) and pBPZpGAL4DBDUw (Addgene 26233), respectively. T2A-p65-Zip^+^ and T2A-Zip^-^-Gal4DBD knock-in constructs were created using two previously described methods: for *upd1-3, Pvf1, Pvf3*, and all *TGFβ* ligands except *gbb*, donor constructs were generated using 1000bp homology arms, amplified via PCR, following the method described in (Ewen-Campen et al. 2020), with an sgRNA targeting a coding exon co-injected. For all other ligand targets, donor constructs were generated using the more efficient “drop-in” method as described in (Ewen-Campen et al. 2023) with the pUC57_Kan_gw_OK2 plasmid backbone from (Kanca et al. 2019; Kanca et al. 2022).

Constructs in the pUC57_Kan_gw_OK2 backbone self-linearize *in vivo* using a synthetic sgRNA that targets either side of the donor, and also encode the gene-specific sgRNA on the same plasmid.

### Transgenic *Drosophila* lines

Knock-in constructs were injected into newly laid eggs expressing germline-restricted Cas9 (*nos::Cas9*). For target genes located on the X, second, or fourth chromosome, we used attP2-*nos::Cas9* (on the third chromosome) as the injection stock, and for any target genes located on the third chromosome, we used attP40-*nos::Cas9* as the injection stock. Injected animals were crossed to an appropriate balancer stock: X chromosome: *FM7*; second chromosome: *CyO*; third chromosome: *TM6b,Tb*; fourth chromosome: *P*{*w[+mC]=ActGFP*}*unc-13[GJ]* (BL9549), and offspring were screened for 3xP3-RFP expression. Approximately four independent transgenic lines were established per target gene, which were then screened for consistent expression. Each line was validated for correct insertion using either PCR genotyping, with a forward primer located 5’ to the left homology arm and a reverse primer located in the T2A sequence of the knock-in, or with GFP expression in a well-characterized domain, or both. All lines in this paper have been submitted to the Bloomington Drosophila Stock Center.

### Antibody staining

Tissues were dissected in PBS, fixed for approximately 30 minutes in 4% paraformaldehyde in PBS, permeabilized in PBST containing 0.3% Triton-X, and stained using standard protocols. Tissues were stained for GFP using Alexa488-coupled anti-GFP (Invitrogen A21311, used at 1:400) and DAPI to visualized nuclei. For larval body wall muscle, a muscle relaxant buffer was used during dissection (5mM EGTA, 5mM MgCl_2_ in PBS) prior to fixation.

Larval muscle was visualized using Alexa555-coupled phalloidin. Sagittal sections of whole adult flies were performed and visualized as described in (Ewen-Campen et al. 2023). Confocal imaging was performed on a Zeiss Axio Observer Z1 with a LSM980 Scan Head, part of the Microscopy Resources on the North Quad (MicRoN) core at Harvard Medical School.

## Acknowledgements

We thank the Microscopy Resources on the North Quad (MicRoN) core at Harvard Medical School for assistance with confocal imaging, and Rich Binari for assistance with fly work, and members of the Perrimon Lab and Drosophila Research Screening Center (DRSC) for valuable feedback. This work was funded by a P41 grant from NIH/NIGMS (5P41GM132087) and from grants from the NIH Office of the Director (5R24OD026435 and 5R24OD031952), and N.P. is an HHMI Investigator. This article is subject to HHMI’s Open Access to Publications policy. HHMI lab heads have previously granted a non-exclusive CC BY 4.0 license to the public and a sublicensable license to HHMI in their research articles. Pursuant to those licenses, the author-accepted manuscript of this article can be made freely available under a CC BY 4.0 license immediately upon publication.

## Literature Cited

Agnes F, Suzanne M, Noselli S. 1999. The Drosophila JNK pathway controls the morphogenesis of imaginal discs during metamorphosis. Development. 126(23):5453–5462. doi:10.1242/dev.126.23.5453.

Augustin H, McGourty K, Steinert JR, Cochemé HM, Adcott J, Cabecinha M, Vincent A, Halff EF, Kittler JT, Boucrot E, et al. 2017. Myostatin-like proteins regulate synaptic function and neuronal morphology. Development. 144(13):2445–2455. doi:10.1242/dev.152975.

Awasaki T, Huang Y, O’Connor MB, Lee T. 2011. Glia instruct developmental neuronal remodeling through TGF-β signaling. Nat Neurosci. 14(7):821–823. doi:10.1038/nn.2833.

Chen Y-CD, Chen Y-C, Rajesh R, Shoji N, Jacy M, Lacin H, Erclik T, Desplan C. 2023. Using single-cell RNA sequencing to generate predictive cell-type-specific split-GAL4 reagents throughout development. Proc Natl Acad Sci. 120(32):e2307451120. doi:10.1073/pnas.2307451120.

Demontis F, Patel VK, Swindell WR, Perrimon N. 2014. Intertissue control of the nucleolus via a myokine-dependent longevity pathway. Cell Reports. 7(5):1481–1494. doi:10.1016/j.celrep.2014.05.001.

Deng M, Wang Y, Zhang L, Yang Y, Huang S, Wang J, Ge H, Ishibashi T, Yan Y. 2019. Single cell transcriptomic landscapes of pattern formation, proliferation and growth in Drosophila wing imaginal discs. Development. 146(18):dev179754. doi:10.1242/dev.179754.

Diao F, Vasudevan D, Heckscher ES, White BH. 2024. Hox gene–specific cellular targeting using split intein Trojan exons. Proc Natl Acad Sci. 121(17):e2317083121. doi:10.1073/pnas.2317083121.

Doherty D, Feger G, Younger-Shepherd S, Jan LY, Jan YN. 1996. Delta is a ventral to dorsal signal complementary to Serrate, another Notch ligand, in Drosophila wing formation. Genes Dev. 10(4):421–434. doi:10.1101/gad.10.4.421.

Everetts NJ, Worley MI, Yasutomi R, Yosef N, Hariharan IK. 2021. Single-cell transcriptomics of the Drosophila wing disc reveals instructive epithelium-to-myoblast interactions. eLife. 10:e61276. doi:10.7554/elife.61276.

Ewen-Campen B, Comyn T, Vogt E, Perrimon N. 2020. No Evidence that Wnt Ligands Are Required for Planar Cell Polarity in Drosophila. Cell Reports. 32(10):108121. doi:10.1016/j.celrep.2020.108121.

Ewen-Campen B, Luan H, Xu J, Singh R, Joshi N, Thakkar T, Berger B, White BH, Perrimon N. 2023. split-intein Gal4 provides intersectional genetic labeling that is repressible by Gal80. Proc Natl Acad Sci United States Am. 120(24):e2304730120. doi:10.1073/pnas.2304730120.

Ewen-Campen B, Mohr SE, Hu Y, Perrimon N. 2017. Accessing the Phenotype Gap: Enabling Systematic Investigation of Paralog Functional Complexity with CRISPR. Developmental Cell. 43(1):6–9. doi:10.1016/j.devcel.2017.09.020.

Housden BE, Perrimon N. 2014. Spatial and temporal organization of signaling pathways. Trends in biochemical sciences. 39(10):457–464. doi:10.1016/j.tibs.2014.07.008.

Hoyer N, Zielke P, Hu C, Petersen M, Sauter K, Scharrenberg R, Peng Y, Kim CC, Han C, Parrish JZ, et al. 2018. Ret and Substrate-Derived TGF-β Maverick Regulate Space-Filling Dendrite Growth in Drosophila Sensory Neurons. Cell Rep. 24(9):2261-2272.e5. doi:10.1016/j.celrep.2018.07.092.

Hu Y, Ewen-Campen B, Comjean A, Rodiger J, Mohr SE, Perrimon N. 2022. Paralog Explorer: A resource for mining information about paralogs in common research organisms. Comput Struct Biotechnol J. 20:6570–6577. doi:10.1016/j.csbj.2022.11.041.

Ishimaru S, Ueda R, Hinohara Y, Ohtani M, Hanafusa H. 2004. PVR plays a critical role via JNK activation in thorax closure during Drosophila metamorphosis. EMBO J. 23(20):3984–3994. doi:10.1038/sj.emboj.7600417.

Jiang H, Patel PH, Kohlmaier A, Grenley MO, McEwen DG, Edgar BA. 2009. Cytokine/Jak/Stat Signaling Mediates Regeneration and Homeostasis in the Drosophila Midgut. Cell. 137(7):1343–1355. doi:10.1016/j.cell.2009.05.014.

Kanca O, Zirin J, Garcia-Marques J, Knight SM, Yang-Zhou D, Amador G, Chung H, Zuo Z, Ma L, He Y, et al. 2019. An efficient CRISPR-based strategy to insert small and large fragments of DNA using short homology arms. Elife. 8:e51539. doi:10.7554/elife.51539.

Kanca O, Zirin J, Hu Y, Tepe B, Dutta D, Lin W-W, Ma L, Ge M, Zuo Z, Liu L-P, et al. 2022. An expanded toolkit for Drosophila gene tagging using synthesized homology donor constructs for CRISPR-mediated homologous recombination. Elife. 11:e76077. doi:10.7554/elife.76077.

Lee JJ, Kessler DPvon, Parks S, Beachy PA. 1992. Secretion and localized transcription suggest a role in positional signaling for products of the segmentation gene hedgehog. Cell. 71(1):33–50. doi:10.1016/0092-8674(92)90264-d.

Lo PCH, Frasch M. 1999. Sequence and expression of myoglianin, a novel Drosophila gene of the TGF-β superfamily. Mech Dev. 86(1–2):171–175. doi:10.1016/s0925-4773(99)00108-2.

Luan H, Diao F, Scott RL, White BH. 2020. The Drosophila Split Gal4 System for Neural Circuit Mapping. Front Neural Circuit. 14:603397. doi:10.3389/fncir.2020.603397.

Luan H, Peabody NC, Vinson CR, White BH. 2006. Refined Spatial Manipulation of Neuronal Function by Combinatorial Restriction of Transgene Expression. Neuron. 52(3):425–436. doi:10.1016/j.neuron.2006.08.028.

Myers L, Perera H, Alvarado MG, Kidd T. 2017. The Drosophila Ret gene functions in the stomatogastric nervous system with the Maverick TGFβ ligand and the Gfrl co-receptor. Development. 145(3):dev157446. doi:10.1242/dev.157446.

Nguyen M, Parker L, Arora K. 2000. Identification of maverick, a novel member of the TGF-β superfamily in Drosophila. Mech Dev. 95(1–2):201–206. doi:10.1016/s0925-4773(00)00338-5.

Nusse R. 2001. An ancient cluster of Wnt paralogues. Trends Genet. 17(8):443. doi:10.1016/s0168-9525(01)02349-6.

Osman D, Buchon N, Chakrabarti S, Huang Y-T, Su W-C, Poidevin M, Tsai Y-C, Lemaitre B. 2012. Autocrine and paracrine unpaired signaling regulate intestinal stem cell maintenance and division. J Cell Sci. 125(24):5944–5949. doi:10.1242/jcs.113100.

Patel A, Wu Y, Han X, Su Y, Maugel T, Shroff H, Roy S. 2022. Cytonemes coordinate asymmetric signaling and organization in the Drosophila muscle progenitor niche. Nat Commun. 13(1):1185. doi:10.1038/s41467-022-28587-z.

Perrimon N, Pitsouli C, Shilo BZ. 2012. Signaling Mechanisms Controlling Cell Fate and Embryonic Patterning. Cold Spring Harbor perspectives in biology. 4(8):a005975–a005975. doi:10.1101/cshperspect.a005975.

Rajan A, Perrimon N. 2012. Drosophila cytokine unpaired 2 regulates physiological homeostasis by remotely controlling insulin secretion. Cell. 151(1):123–137. doi:10.1016/j.cell.2012.08.019.

Rosin D, Schejter E, Volk T, Shilo B-Z. 2004. Apical accumulation of the Drosophila PDGF/VEGF receptor ligands provides a mechanism for triggering localized actin polymerization. Development (Cambridge, England). 131(9):1939–1948. doi:10.1242/dev.01101.

Simcox AA, Grumbling G, Schnepp B, Bennington-Mathias C, Hersperger E, Shearn A. 1996. Molecular, Phenotypic, and Expression Analysis ofvein,a Gene Required for Growth of the Drosophila Wing Disc. Dev Biol. 177(2):475–489. doi:10.1006/dbio.1996.0179.

Tabata T, Eaton S, Kornberg TB. 1992. The Drosophila hedgehog gene is expressed specifically in posterior compartment cells and is a target of engrailed regulation. Genes Dev. 6(12b):2635–2645. doi:10.1101/gad.6.12b.2635.

Tirian L, Dickson BJ. 2017. The VT GAL4, LexA, and split-GAL4 driver line collections for targeted expression in the Drosophila nervous system. Biorxiv.:198648. doi:10.1101/198648.

Upadhyay A, Moss-Taylor L, Kim M-J, Ghosh AC, O’Connor MB. 2017. TGF-β Family Signaling in Drosophila. Cold Spring Harb Perspect Biol. 9(9):a022152. doi:10.1101/cshperspect.a022152.

Upadhyay A, Peterson AJ, Kim M-J, O’Connor MB. 2020. Muscle-derived Myoglianin regulates Drosophila imaginal disc growth. eLife. 9:e51710. doi:10.7554/elife.51710.

Wharton KA, Ray RP, Gelbart WM. 1993. An activity gradient of decapentaplegic is necessary for the specification of dorsal pattern elements in the Drosophila embryo. Development. 117(2):807–822. doi:10.1242/dev.117.2.807.

Worley MI, Everetts NJ, Yasutomi R, Chang RJ, Saretha S, Yosef N, Hariharan IK. 2022. Ets21C sustains a pro-regenerative transcriptional program in blastema cells of Drosophila imaginal discs. Curr Biol. 32(15):3350-3364.e6. doi:10.1016/j.cub.2022.06.040.

Yan S-J, Gu Y, Li WX, Fleming RJ. 2003. Multiple signaling pathways and a selector protein sequentially regulate Drosophila wing development. Development. 131(2):285–298. doi:10.1242/dev.00934.

Zee M van der, Fonseca RNda, Roth S. 2008. TGFbeta signaling in Tribolium: vertebrate-like components in a beetle. Development Genes and Evolution. 218(3–4):203–213. doi:10.1007/s00427-007-0179-7.

